# PALAEOROBOTICS UNCOVERS DEVASTATING MAMMALIAN TAIL STRIKE DYNAMICS

**DOI:** 10.64898/2026.04.21.718724

**Authors:** Kévin Le Verger, Federico Allione, Bingcheng Wang, Bernhard Weisse, Auke J. Ijspeert, Marcelo R. Sánchez-Villagra, Torsten M. Scheyer, Ardian Jusufi

## Abstract

Extant animals occupy only a fraction of evolutionary design space, leaving many extinct morphologies without living analogues for functional testing. We combined comparative anatomy of 32 glyptodont species with a life-sized palaeorobotic tail rig to determine strike severity in club-tailed giant from the Pleistocene megafauna *Doedicurus*. The robophysical model, informed by fossil geometry and inertia, matched simulated fossil-tail collision momentum (98-105%) and generated peak forces of 81.1 kN at 6.4 m·s⁻¹, with an impulse of 427 N·s, far exceeding equine extremity fracture forces. Peak force scaled near-linearly with strike velocity, implying ∼183 kN at projected top speed of 15 m·s⁻¹. Measurements suggest *Doedicurus* delivered high-severity strikes, supporting antipredator defence as primary selective driver of extreme tail weaponization, and establish palaeorobotic validation of extinct animal biomechanics.

## Main Text

The fossil record demonstrates that modern morphological diversity is only a subset of what has evolved in the geological past. Thus, much of the past diversity captures regions of morphological space that are no longer occupied by living organisms (*1*). Understanding the evolution and function of such structures requires the integration of approaches (*2–4*). Numerous fossil examples document evolutionary arms races driven by prey-predator interactions (*5–7*), leading to the coevolution and escalation of traits related to predation and antipredator defences. Among the most prominent of these is bony armour, as observed in placoderms (*8*), many reptiles (*9*), including ankylosaur dinosaurs, and mammals such as armadillos (*10*), ranging from scattered osteoderms to continuous protecting coverings resembling knight’s chainmail or full plate armour. In several armoured herbivorous clades, the tail further exhibits morphological specialisations consistent with its use as a weapon (*11*). Such tail weaponisation may function either as a defensive response to predation or in intraspecific combat, among conspecifics (*11*). Distinguishing between these selective pressures remains challenging, and the adaptive significance of weaponised tails is increasingly debated (*12*). In some armoured prey, several studies favour a higher impact from conspecific fighting than predation pressure as a driver for tail weaponisation without yet experimentally determining it (*11, 13, 14*). Analogous to the ankylosaur, a conspecific-driven hypothesis is also preferred for the gigantic glyptodonts based on the presence of osteoderms with traces of deformation (*10*), caudal vertebral pathologies (*15, 16*), and several studies on their tail function (3, 17, 18). Glyptodonts were gigantic herbivorous armadillos that roamed the Americas, with an evolutionary history marked by repeated adaptive acquisitions towards an armoured body and a weaponised tail (*19*). Considering their pronounced tail anatomical diversity and the advent of the Great American Biotic Interchange as a source of new large-bodied predators (*20*), such as sabre-toothed cats, like *Smilodon* (with the largest individuals exceeding 400 kg (*21*)), and the gigantic bear *Arctotherium* (with body masses exceeding 1,700 kg (*22*)), glyptodont tails represent an ideal model for testing the potential interplay between predatory and intraspecific pressure. In these extinct animals, the tail is inferred from both its morphology and overall body design to have constituted the only active defensive structure. In encounters with such large, agile megapredators, effective tail strikes would therefore have needed to be decisive, either incapacitating or deterring the attacker, before the animal could rely solely on its carapace. This functional interpretation requires validation through biomechanical analyses. Previous estimates of glyptodont tail movement and impact forces have relied on theoretical mathematical calculations (*3, 17*), which have yet to be validated in practice. Following an anatomical assessment of 32 glyptodont species, covering 19 million years of evolution, we used palaeorobotics on the most heavily weaponised species to test the force on impact of tail strike dynamics. We determined the damage that this weapon could have caused to a conspecific or predator, assuming that an excessively lethal tail strike would have resulted preferentially from anti-predator adaptation. We show that one glyptodont lineage (*i.e.*, Hoplophorinae) experienced a selection pressure towards tail weaponry because of evolutionary radiation at the end of the Miocene, supposing the strong involvement of intraspecific interactions. One clade emerging from this diversity peak has yielded the most heavily weaponised glyptodont during the Pleistocene, the two tons club-tailed *Doedicurus clavicaudatus*. The dynamic palaeorobotic model of which reveals a devastating tail strike at a speed of 6.4 m·s⁻¹ that yielded an impact force of 81.1 kN, exceeding equine bone fracture thresholds (*23, 24*). Thus, the projected top speed of 15 m·s⁻¹ would generate 183 kN, which is comparable to car crash forces experienced (*4*). Consequently, we argue that the tail of *Doedicurus* is primarily optimised for deterring or incapacitating Pleistocene mega-predators and, only incidentally, for confronting a conspecific. Increasing predation pressure would have acted as a catalyst in the selection towards tail weaponisation in this mammal.

### Tail diversity and defence index

Defined as a succession of caudal rings composed of osteoderms, ending in either a small tubercle or a long caudal tube (Fig. 1), the caudal armour of glyptodonts has no equivalent in the animal kingdom. Anatomical differences can be examined to determine how the tail is associated with a defence type. To investigate this link, we examined 19 of the 25 morphological characters derived from caudal armour in the most recent time-calibrated glyptodont phylogeny (*25*). By converting phylogenetic characters into ordered anatomical defence features, we can code anatomical variations in the form of defence scores, from the most protective morphological traits to the most offensive ones (fig. S1 and data S1). The states derived from the phylogenetic characters are thus transformed into ordered scores for a given anatomical feature using the following three rationales: (1) a long caudal tube is considered more optimal for striking than a terminal tubercle; (2) an optimised tail for striking is supposed to show less ornamentation for improved robustness and high vascularity for the blood supply; (3) any complexifications of the tail shape and surface (excluding ornamentation) of the distal end is here considered as weaponisation. For a given taxon, the sum of the scores divided by the number of coded features gives a defence value allowing the classification of glyptodonts along a spectrum of defence (see Materials and Methods), which we refer to a defence index herein. As a result, our defence index ranges from 1.1, indicating a highly protective mode, to 4.1, meaning a highly offensive mode (data S2). Among 32 glyptodont species (fig. S2), four major morphotypes of caudal armour can be recognised (Fig. 1). The first morphotype, the most protective mode (defence index < 2.0) is represented by glyptodonts with a terminal tubercle, all belonging to the Glyptodontinae (7/32). In these glyptodonts, the tail is relatively massive, composed mainly of unfused caudal rings, and ends distally with a terminal tubercle. The second morphotype (2.0 < defence index < 3.0) is the most widespread anatomy (17/32), corresponding to a simple ornamented caudal tube with no apparent signs of a more complex anatomy, showing weaponisation. In contrast, the next two morphotypes show the acquisition of weaponry (*e.g.*, lateral depression and dorsoventral flattening). Among the most offensive modes, the third morphotype (3.0 < defence index < 4.0) includes members of the Doedicurini and Hoplophorini tribes (7/32). The Panochthini are worthy representatives with their ‘Viking sword’ tail, so named because of its straight, dorso-ventrally flattened caudal tube and the presence of a multitude of depressions with a small spine-like structure at their centre (*18*). Two of the multiple depressions are much larger in the latest morphotype (defence index > 4.0), only represented by the club-tailed *Doedicurus clavicaudatus* (1/32). In this glyptodont, the caudal tube has lost a large part of its ornamentation through osteoderm fusions, and exhibit a dorsoventral flattening, the presence of huge depressions, and a distal part ending in a sledge-hammer shape. Along this spectrum, Glyptodontinae, and *Doedicurus* in particular, represent extreme morphologies within the broader diversity of glyptodonts, making *Doedicurus* a compelling model for tail strike investigation.

**Fig. 1.**
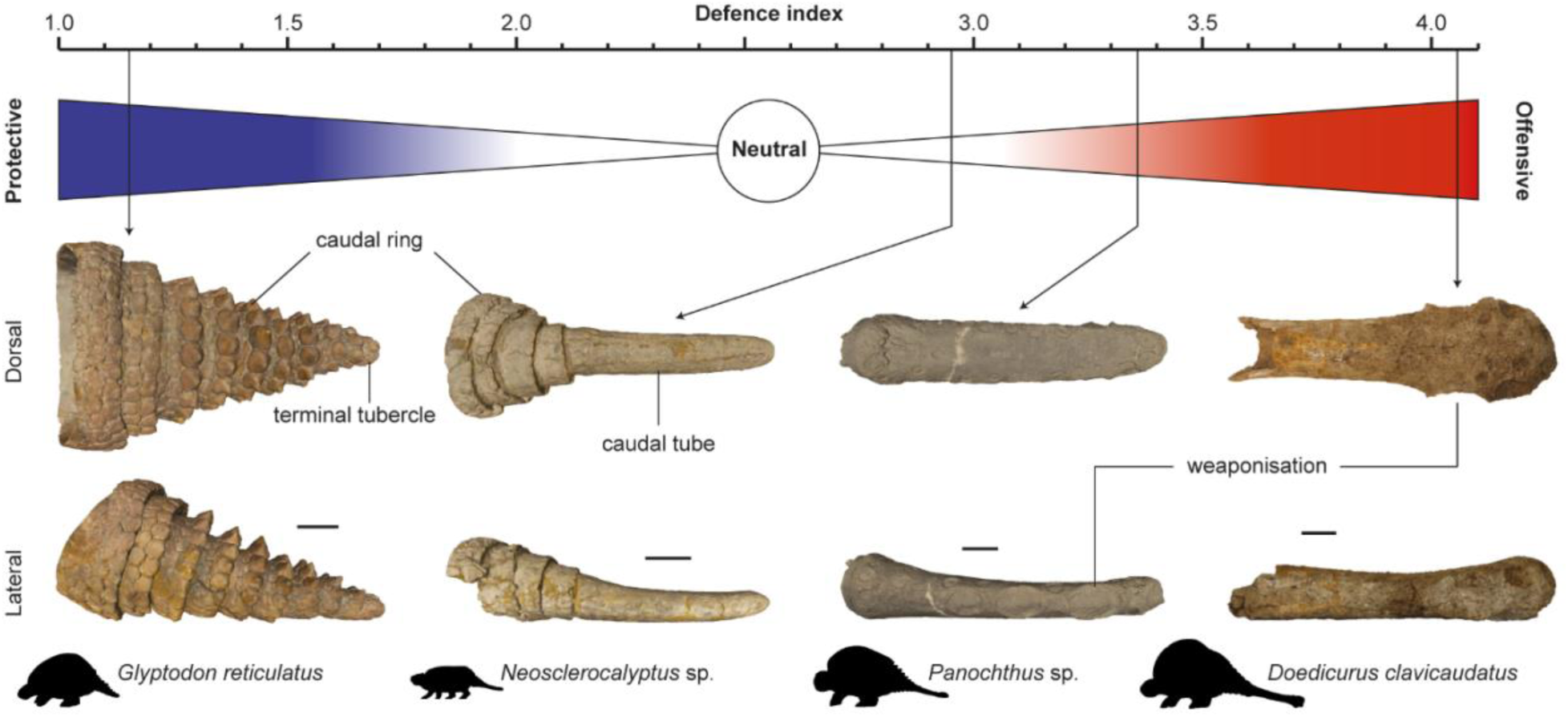
The defence index derived from the glyptodont tail anatomy and its four morphotypes. The four types of tail, from the most protective to the most offensive, are shown with key anatomical indications. From left to right, the fossils illustrated are MACN-Pv 1780, MACN-Pv 9630, MACN-Pv 14998 (Museo Argentino de Ciencias Naturales Bernardino Rivadavia, Paleovertebrados, Buenos Aires, Argentina) and PIMUZ A/V 459 (Natural History Museum, University of Zurich, Switzerland). The silhouettes are handmade. Scale bars, 10 cm.

### Impact dynamics experiments

The speed of movement at the tip of the tail (15 m·s⁻¹) and the energy required to break the carapace of a conspecific (1.4 to 6 kJ, thus an impact force of 140 to 600 kN considering a displacement of 1 cm) were estimated by Alexander *et al*. (*3*) based on theoretical mathematical approaches with numerous assumptions. Since then, this study has become a benchmark and no new estimates have been attempted, although the tail functionality (*17, 18*), the carapace robustness (*26*), and evolutionary convergence with ankylosaurs (*27*) have been questioned. Because *Doedicurus* appears to be a unique case of an ultra-offensive tail in accordance with our defence index (Fig. 1), we focused our robotic experimentation on the tail of this species. To determine the forces developed on impact in a simplified dynamic test rig, we built a life-sized dynamic palaeorobotic physical model to approximate the salient mechanical features of the glyptodont fossil tail (Fig. 2).

**Fig. 2.**
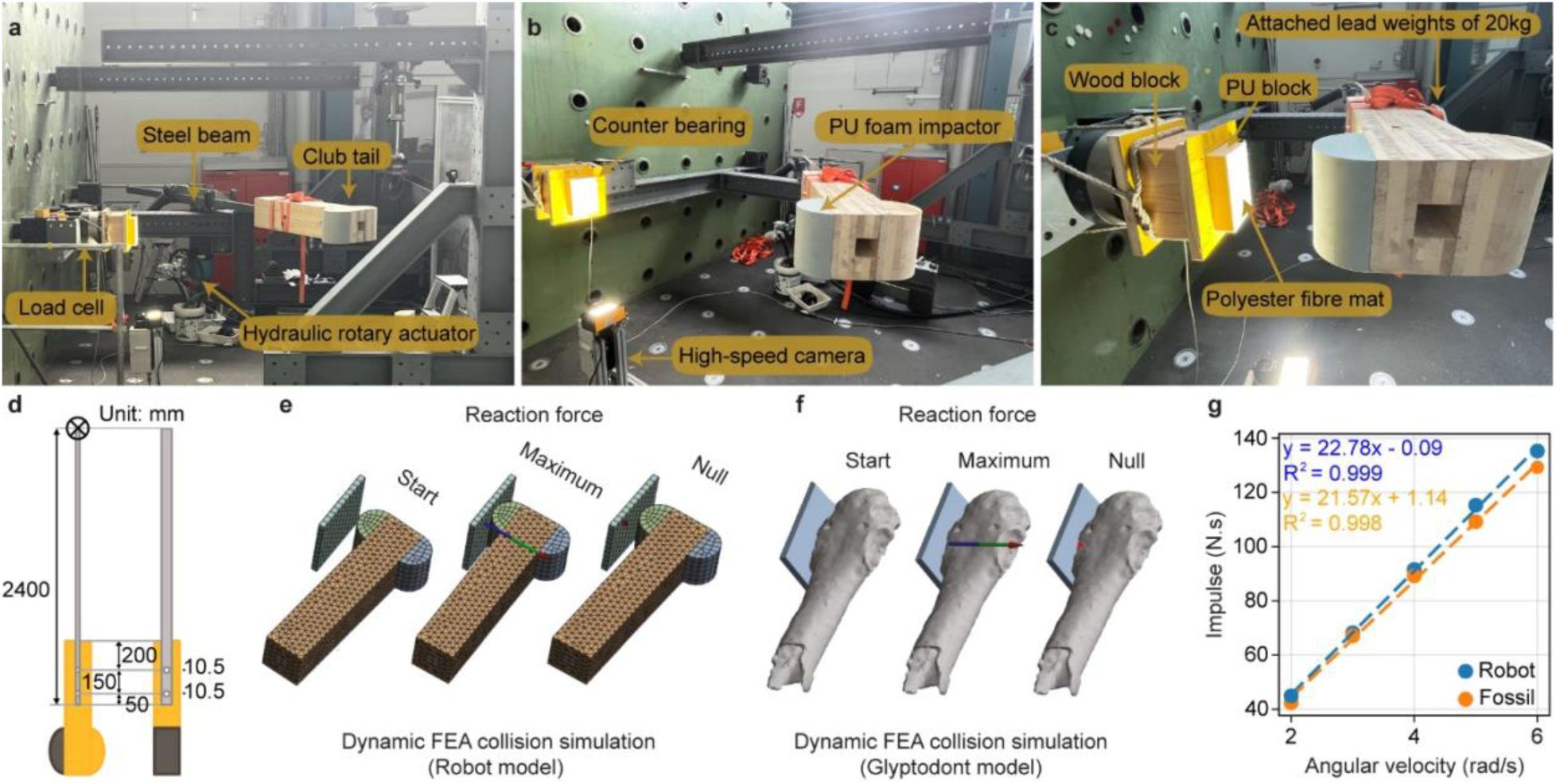
Dynamic experimental validation of glyptodont tail strikes informed by analytical modelling. Life-sized palaeorobotic strike rig with hydraulic motor accelerating large tail, emulating fossil mass distribution and inertia in side view (**A-C**). Bone density material representing the glyptodont tail club (SYNBONE) was mounted via wooden elements to the metal beam, facing the target force platform. Dimensions of 2.4 m long tail and distal club in top and side views (**D**). The dynamic FEA collision simulation of the robophysical model (**E**) and the analytical glyptodont model (**F**) are displayed. Comparison of impulse generation in the glyptodont simulation and the robotic experiment (**G**). Both systems were initialised with identical angular velocities, with all external torque removed at the point of impact to ensure a purely inertial collision.

Given the high proximal flexibility due to several segment joints and the long rigid distal part formed by the caudal tube at the base, similar to a joystick, the simplification of the motion for the test is strongly supported by previous studies, while the dynamic similarity of the salient features of the tail strike mechanics relevant to the hypothesis in question is maintained. For instance, the horizontal motion has already been proposed as the optimal use for tail striking (*18*) and the centre of percussion was determined at the level of the largest distal depressions (*17*). Moreover, numerous additional factors (*e.g*., weight, dimensions, density) were considered in the device development of the robophysical model, design and motion profiles (see Materials and Methods). To demonstrate that a paleorobotic physical model with appropriate inertia can accelerate the high-density tail club (SYNBONE) to reconstruct the impact performance of fossil tail clubs, we analysed the impact dynamics of the biomimetic palaeorobotic tail and the glyptodont fossil tail under identical conditions in Finite Element Analysis (FEA). The dynamic palaeorobotic model achieved impact momentum values that were between 98% and 105% of those of the simulated glyptodont fossil tail at five impact velocities tested (Fig. 2). This close correlation indicates that the palaeorobotic physical model of the tail effectively approximates the mass distribution and centre of mass position of the glyptodont tail, resulting in inertial characteristics and momentum, such that the impact dynamics measured (Fig. 3) provide insights for how glyptodonts may have utilised their tails with respect to the biomechanics.

**Fig. 3.**
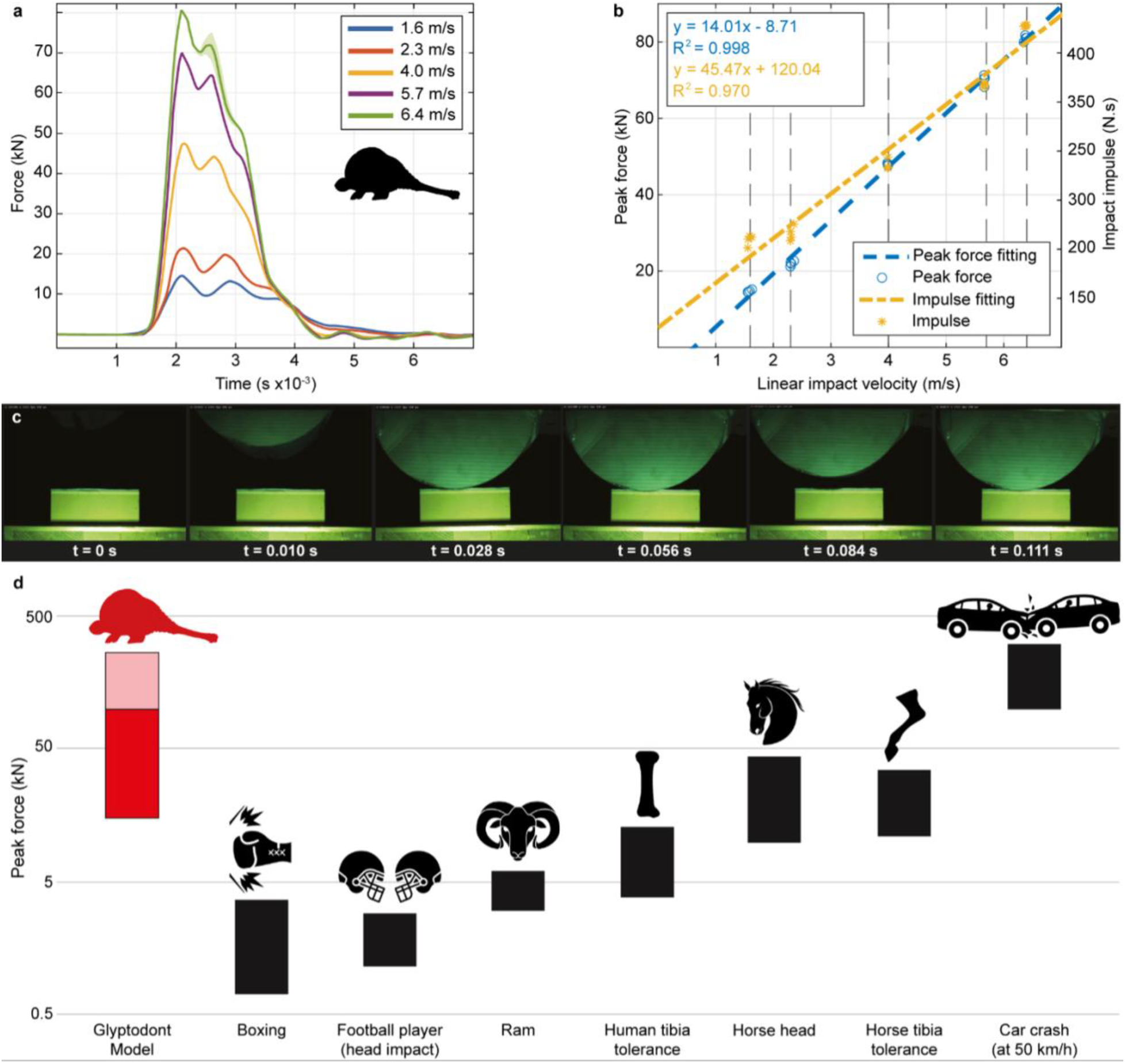
**Impact dynamics experiments of life-sized glyptodont robophysical model tail strikes powered by a hydraulic motor**. (**A**) Force-time histories throughout tail club strikes at speeds ranging from 1.6 to 6.4 m·s⁻¹ onto instrumentalised force platform. (**B**) Illustration of peak forces, and impact moment, respectively, determined at 5 collision speeds and their fitting results. (**C**) Time-series image sequence reveals close up of the tail club approaching the target contact zone and impact during strike events. (**D**) Comparison of peak forces for glyptodont tail club impact with other collisions in extant animals, including ram head-butting and equine skull/tibia fractures, differences relative to today curve obtained based on benthic foraminifera δ^18^O values from Westerhold *et al*. (*37*) with smoothing over 20 kyr (grey) and 1 Myr (red). Major climatic and environmental events follow the indications of Westerhold *et al*. (*37*) and Núñez-Blasco *et al*. (*25*). Specific diversity of the four main groups of predators is derived from Prevosti *et al*. (*20*). Phylogeny is adapted from Núñez-Blasco *et al*. (*25*). Colour coding represents the value of the defence index of the 32 glyptodont species investigated (fig. S2), also reconstructed at each node by maximum likelihood following Brownian motion. Body mass data correspond to a compilation of previous research, details of which are given in Materials and Methods. All these variables are plotted against the geological time scale (GTS2012). Illustrated tails are not to scale. Abbreviations: Calab., Calabrian; Exp., expansion; Gel., Gelasian; Mean temp. diff. to today, mean temperature difference to today; Mid., Middle Pleistocene; Piac., Piacenzian. Symbol: *, Upper Pleistocene. All silhouettes are handmade. as well as human biomechanics, including boxing, American football athletes head collisions, and car crash biomechanics (see Methods). Except for the *Doedicurus* silhouette, credits from left to right in (D) refer to Lê Khắc Bảo Thoại, Nawicon, Damian Patrignani, iconfield, yogi rista, NHA, and Adriana Danaila, from the Noun Project.

By approximating the geometry and mass distribution of the glyptodont tail through analytical and physical modelling, we were able to create a dynamic palaeorobotic physical model that closely approximates the salient features of its natural counterpart and exhibits comparable mechanical conditions on impact. The force-time history measurements reveal the impact forces generated by the glyptodont biomimetic tail at different linear impact velocities of 1.6 to 6.4 m·s⁻¹ (Fig. 3).

As the velocity increases, the peak impact force significantly increases. At 1.6 m·s⁻¹, the peak force reaches approximately 15 kN, with an impulse of 201.3 N·s. At 4.0 m·s⁻¹, we measured a peak force exceeding 48.3 kN, with an impulse of 293.8 N·s. At the highest speed our test rig was still safe to operate, that is 6.4 m·s⁻¹, the peak force reached 81.1 kN, with an impulse of 427.0 N·s. Force-time history profiles show a sharp rise to the peak force followed by a gradual decline. Higher velocities result in not only higher peak forces but also more complex force profiles with multiple peaks and broader force durations. The time series image sequence corresponding to the close-up view of the collision can be seen in Fig. 3C and the corresponding high-speed video recorded at 1000 Hz is available for viewing (movie S1). The scatter plot in Fig. 3B correlates peak impact force with linear impact velocity. The trend line indicates that as the impact velocity increases, the peak impact force rises in a near-linear fashion. This linear relationship suggests that the palaeorobotic tail’s impact force can be reliably predicted based on the impact velocity, with higher velocities producing proportionately higher impact forces. Consequently, considering the speed of 15 m·s⁻¹ proposed by Alexander *et al*. (*3*), our estimates result in an impact force of 183 kN, the equivalent of forces experienced by humans during car crashes (Fig. 3D), making the *Doedicurus* tail a formidable weapon.

## Discussion

Anatomical structures found in extinct animals have raised a plethora of questions about the paths taken during the course of evolution. With their unique, yet highly diverse caudal armour (*19*), glyptodonts are worthy representatives of those vestiges of the biological past for which the tail function has remained a mystery. Over the last 40 years, interpretations of glyptodont tail weaponisation have shifted from a predator-driven hypothesis, where a heavy carapace and weaponised tail evolved primarily in response to predation pressure, towards a conspecific-driven hypothesis emphasising intraspecific combat (*3, 17, 27*), leading to the emergence of a narrative centred on glyptodont intraspecific warfare (*28*). One of the main studies supporting the conspecific hypothesis is the theoretical mathematical approach proposed by Alexander *et al*. (*3*), which suggested that the range attributed to the energy developed by the tail to perform a strike was likely equivalent to the potential energy to fracture the carapace. However, this inference was based on indirect estimates, and the actual impact forces generated during a tail strike remained to be measured. Additional support for the conspecific hypothesis has been drawn from strong phenotypic convergence between glyptodonts and ankylosaurs, as highlighted by Arbour & Zanno (*27*). In ankylosaurs, extensive work by Arbour and colleagues has argued that the complex morphology, expensive physiology, and high taxonomic variability of tail clubs are more consistent with sexually selected weapons than with antipredator adaptations (*11, 14, 27*). By analogy, this convergence has been interpreted as favouring a similar conspecific and sexually selected function for glyptodont tail weaponisation (*27*). By contrast, ankylosaurs relied on a distinct structural strategy for tail weaponisation, involving proximodistally arranged ossified tendons that produced a highly rigid club (*29*). Impact forces for ankylosaur tail strikes have been inferred from biomechanical models and are substantially lower than those measured experimentally here for *Doedicurus*, even though these approaches are not directly comparable^29^. Although the image of two glyptodonts fighting like two knights may have been seductive, the empirical evidence supporting this scenario remains fragile because traces of secondary healing of the dorsal carapace are rare and tough to diagnose (*10*). Moreover, almost no agonistic behaviour is known in extant armadillos, only minor variations in the armour of Glyptodontinae have been identified as potential sexual dimorphisms (*30*), and there was no significant difference in the external anatomy of the armour plan despite a wide variation in tail weaponisation (*19*). In other words, the role of tail weaponry, and therefore of the selective mechanisms behind the acquisition of these extraordinary biological traits, remained open for debate.

Our results revealed that the weaponised tail as an offensive defence is a relatively rare feature within glyptodonts, finding its optimum in their giant representative, *Doedicurus*. Thanks to palaeorobotics, an actuated life-sized physical model of the tail imitating the salient mechanical features of its gigantic natural counterpart enabled the force on impact of a weaponised glyptodont tail to be measured for the first time. Impact dynamics experiments revealed the force on impact of a *Doedicurus* tail strike to be devastating, reaching 81.1 kN at a speed of 6.4 m·s⁻¹, which surpasses forces measured during equine skull and tibiae fractures (*23, 24*), while 183 kN are projected at the maximal speed (*3*) of 15 m·s⁻¹, which falls within the range found in car crashes (*4*) (Fig. 3). Bearing in mind that our approach is a simplified one, offering a conservative estimate, the caudal armour of *Doedicurus* could have been an even deadlier weapon with potentially even more devastating motion profiles, amplitudes, inertia from consecutive tail oscillations, and the probable presence of keratinous spines, comparable to rhinoceros’ horn features (*3, 17*).

In this study, we focus on peak reaction force because it is directly measured in our experimental set up (hydraulic actuator test rig with 10 kHz load cell) and provides a simple benchmark to compare strike severity across velocities in a controlled, repeatable way, including the measured force–time histories up to 81.1 kN at 6.4 m s⁻¹ and the increasingly complex, multi-peaked pulses at higher speeds. Nevertheless, we are fully aware of the limitations of our study, including that impact dynamics in systems in motion cannot be reduced to a single scalar. Impact dynamics indeed depends on many factors, including the contact area and stress distribution, the full force-time history and contact duration τ, the impulse 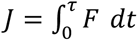, and the work/energy exchange 𝑊 = 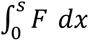, with 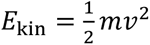 partitioned into deformation, heating, fracture, and recoil. These materials and mechanics relationships are explicit in the standard scalings 𝐹^ˉ^ ≈ Δ𝑝/𝜏 ≈ 𝑚𝑣/𝜏 and 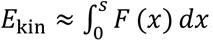, which highlight that the same peak force can arise from very different combinations of τ and stopping distances. Tail strikes are energy-transfer events, where kinetic energy is imparted from the accelerated tail tip to the impacted body (predator and/or conspecific), while some fraction is retained within the glyptodont tail club system as rebound and internal vibrations. Interpreting ecological consequences therefore also requires attenuation, meaning how joint compliance, soft tissues, and stiffness gradients act as a load-absorbing region (*i.e.*, engineering analogue of crumple zone) that reshapes the pulse and limits proximal load transmission. Our peak force comparisons should therefore be read as an externally measurable envelope under controlled conditions rather than a complete description of internal load paths. Holistic comparative analysis across extant species and human collision events, as attempted in Fig. 3D, are inherently challenging because many studies do not report the full set of variables needed to map peak force onto impulse, energy dissipation, contact mechanics, and internal dynamics, including the fact that, for extinct systems, the boundary conditions and tissue-level properties are especially hard to constrain.

Relatively high forces were previously estimated for fracturing the carapace of a glyptodont (ranging from 140 to 600 kN assuming displacement of 1 cm for energy dissipation (*3*)), particularly in comparison with the resistance to impact forces known in many other herbivores, as exemplified in the case of head-butting (*31*). Nevertheless, the values we obtained for the impact forces of a tail strike do fall within this range. Alexander *et al*. (*3*) compared values for human and goat skulls with glyptodont carapaces, which could be seen as a substantive underestimate if we consider the relative thickness, osteoderm-connectivity, and rigidity of the latter, as described by the authors themselves. More recently, Du Plessis *et al*. (*26*) provided evidence that the histological composition of glyptodont osteoderms could approach optima in combination of strength and high energy absorption. Having tested the energy absorption of a glyptodont osteoderm configuration with different porosities and, despite the limitations of their mechanical tests (up to 30 kN and 100 J), their results revealed shock absorption capacities in much more plausible force and energy ranges. Du Plessis *et al*. (*26*) also demonstrated that excessive porosity limited this structural advantage for energy absorption. The osteoderms of the dorsal carapace of *Doedicurus* show a relatively high porosity (37.85%; fig. S3), implying a resistance substantially lower than that proposed by Alexander *et al*. (*3*) and far from withstanding the optimal impact of a conspecifics′ tail clubs. More importantly, the osteoderms of *Doedicurus* have large canals for blood supply (fig. S3), which would have tended to reduce the resistance of an osteoderm to impact even more. Another structural advantage could lie in the morphology of the carapace with a dome-like protection provided by several interconnected osteoderms. Perhaps such structural organisation would also increase shock absorption, a promising prospect for bioinspired design and the conception of glyptodont armour. An obvious follow-up study to ours would be to repeat the tests of Du Plessis *et al*. (*26*) with greater force and energy applied, to experimentally determine the structural integrity of the entire carapace under dynamic impact. For the moment, the aforementioned evidence suggests that the impact force of a *Doedicurus* maximum tail strike is much higher than the absorption capacity of the dorsal carapace. Although the maximum force of the tail is tremendous, likely lethal to any adversary, this finding does not necessarily conflict with the idea of agonistic behaviour between glyptodonts. Intraspecific fights are more likely than interspecific fights within glyptodonts given the size differences (*i.e.*, from hundreds of kilograms to tons), although the same impact energy can be obtained with different solutions (fast and light vs. slow and heavy), and the usually relatively non-antagonistic interactions between large herbivores of different species. More particularly in the case of non-predators, when two individuals of the same species engage in combat, the aim appears to be to intimidate the opponent rather than to kill (*32*). Imagining that glyptodonts did not use the maximum power of the tail during their confrontation to limit severe lesions on the dorsal carapace and caudal vertebrae is thus parsimonious. The tail could therefore clearly have been used against a predator as well as a conspecific, but intraspecific interactions alone may not explain the optimisation in tail weaponry seen in *Doedicurus*.

To gain a better understanding of the history behind the optimisation of *Doedicurus* tail weaponry in the light of our results, we placed our investigation in an evolutionary context. Caudal armour evolution in glyptodonts took place in a period of 19 million years in which environments were disrupted by biotic and abiotic factors (*33–35*). To examine the evolution of the defence spectrum in relation to these major upheavals, we reconstructed the ancestral state of the tails being utilised for defence using maximum likelihood under a Brownian motion on the time-calibrated morphological phylogeny of glyptodonts (*25, 36*) (*i.e.*, defence index (y) vs. time (x)). We then temporally calibrated this reconstruction with first appearance data (*25*), and added consideration of the variation in body size (see Materials and Methods), global climatic data (*37*) and the specific diversity of predators in South America (*20*) (Fig. 4 and data S2).

**Fig. 4.**
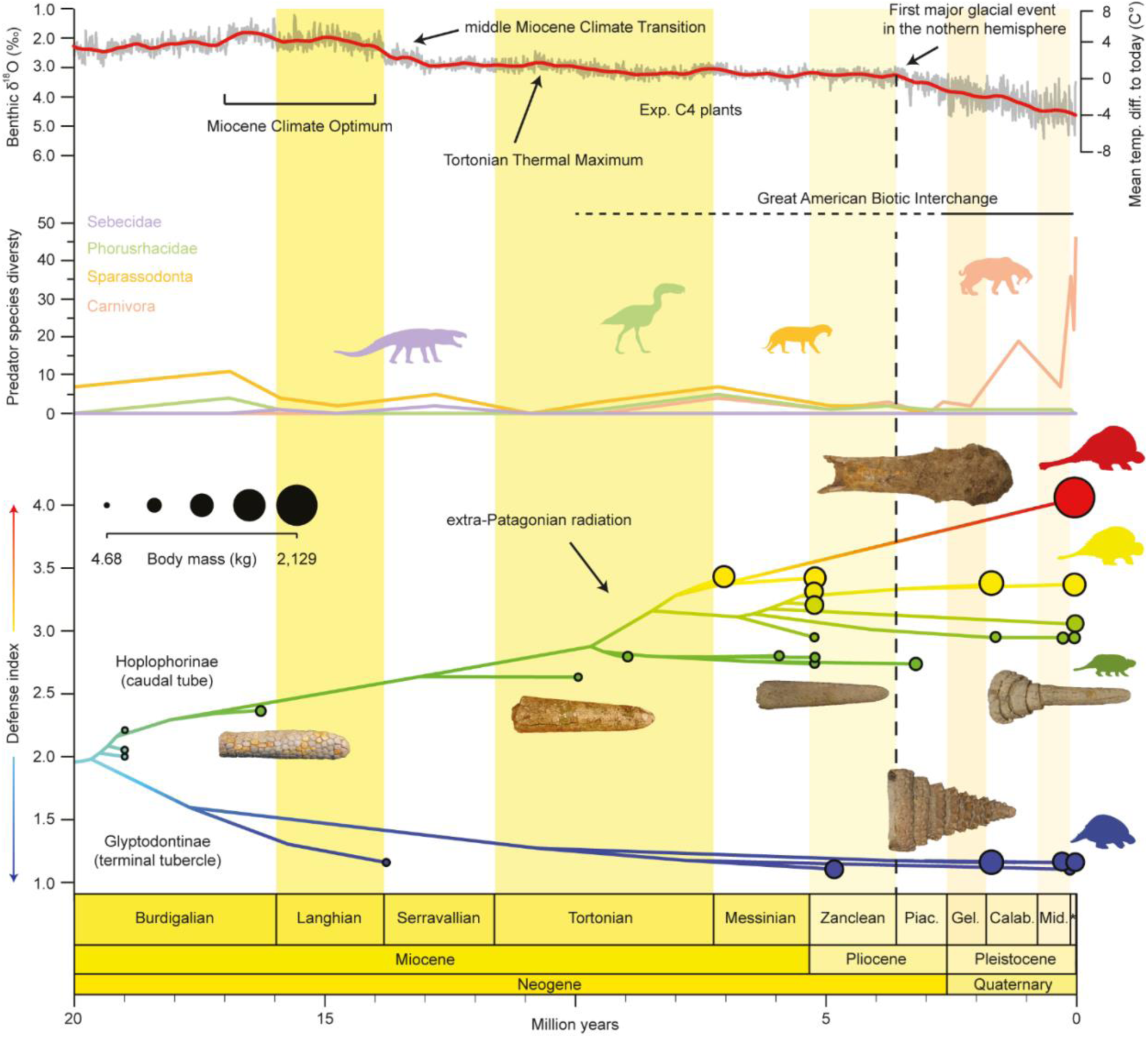
Evolution of glyptodont tail defence index in relation to paleoclimate, South American predator diversity and body mass during the Neogene and Quaternary. Mean temperature differences relative to today curve obtained based on benthic foraminifera δ^18^O values from Westerhold *et al*. (*37*) with smoothing over 20 kyr (grey) and 1 Myr (red). Major climatic and environmental events follow the indications of Westerhold *et al*. (*37*) and Núñez-Blasco *et al*. (*25*). Specific diversity of the four main groups of predators is derived from Prevosti *et al*. (*20*). Phylogeny is adapted from Núñez-Blasco *et al*. (*25*). Colour coding represents the value of the defence index of the 32 glyptodont species investigated (fig. S2), also reconstructed at each node by maximum likelihood following Brownian motion. Body mass data correspond to a compilation of previous research, details of which are given in Materials and Methods. All these variables are plotted against the geological time scale (GTS2012). Illustrated tails are not to scale. Abbreviations: Calab., Calabrian; Exp., expansion; Gel., Gelasian; Mean temp. diff. to today, mean temperature difference to today; Mid., Middle Pleistocene; Piac., Piacenzian. Symbol: *, Upper Pleistocene. All silhouettes are handmade.

At the beginning of the Miocene, glyptodonts underwent an ancestral dichotomy with the divergence of two distinct lineages, one with a caudal tube, the austral clade, and the other with a terminal tubercle, the Glyptodontinae. At this point, the defence index only reflects protective modes of defence for the use of caudal armour. The two lineages remained poorly diversified during the gradual cooling that occurred in the Miocene. However, the austral clade underwent an evolutionary radiation during the Late Miocene, called the extra-Patagonian radiation (*38*), accompanied by an increase in the defence index in several new glyptodont species (Fig. 4). This radiation could be explained by the increase in arid conditions and the proliferation of C_4_ plants in South America due to climatic cooling during the Neogene (*37–41*), conditions that are favourable to glyptodonts which are browse/grass-dominated mixed-feeders (*42, 43*). No abiotic factor alone accounts for the increase in the defence index. During late Miocene, climatic conditions remained relatively stable and, although predation pressure from terror birds and marsupial predators increased, the rise was moderate compared with the early Miocene (*20*), particularly following the disappearance of terrestrial sebecosuchian crocodylomorphs. A more plausible explanation is an increase in intra- and interspecific competition in an ever more open environment, consistent with a probable intensification of sexual selection. This period also coincided with a marked increase in body size, with several glyptodont lineages reaching several hundred kilograms, and with the late Miocene radiation that gave rise to most Pleistocene glyptodont tribes. A major shift occurred during the Pliocene and Pleistocene. Global cooling following the Pliocene Climatic Optimum promoted open habitats and drove the gigantism characteristic of Pleistocene megafauna (*44*), with some glyptodonts, including *Doedicurus*, reaching one to two tonnes (*45, 46*). Concurrently, the formation of the Isthmus of Panama intensified the Great American Biotic Interchange, leading to a sharp increase in predator diversity with the arrival of large carnivorans in South America (*35*). Glyptodonts were thus confronted with novel megapredators, including sabre-toothed cats, larges canids and bears (*47*). Functionally comparable to heavily armoured tanks, glyptodonts possessed a continuous carapace covering most of the body (*48*), but unlike some armadillos or turtles, they were unable to curl up or retract their appendages (*49–51*), leaving vulnerable regions exposed. Under these conditions, an active defensive weapon would have been crucial. Unlike intraspecific combat, antipredator defence favours maximal damage rather than intimidation, consistent with strong selection on tail weapon performance. During the Pleistocene, the defence index reached its highest values in *Doedicurus*, which emerges as a clear outlier in caudal armour. Several localities document the coexistence of *Doedicurus* with some of the largest and most agile predators of the time (*52, 53*). Unable to outrun such attackers (*54, 55*), glyptodonts likely relied on active defence or on limiting access to vulnerable body regions, particularly in encounters involving juveniles, senescent or injured individuals. This pressure would have been exacerbated if predators such as sabre-toothed cats or South American canids engaged in social hunting (*56*). Although gregarious behaviour is unknown in extant armadillos, it is common among large herbivores (*57, 58*) and cannot be excluded in glyptodonts, potentially allowing groups to confront predators collectively, albeit using tail strikes rather than tusks or horns. In summary, the evolutionary radiation of glyptodonts during the Late Miocene likely promoted the diversification of tail morphotypes. In the context of increasing glyptodont abundance, traits associated with offensive tail use may initially have been favoured in intraspecific interactions, potentially driven by sexual selection. By contrast, the pronounced climatic cooling beginning in the Pliocene, coupled with the intensification of predation pressure following the Great American Biotic Interchange, may have acted as a catalyst for directional selection towards extreme weaponised morphologies. In this later context, tail weaponry may have been increasingly shaped by antipredator defence rather than by conspecific combat. Although glyptodont tails may have served additional functions, such as balance or environmental interaction, our dynamic palaeorobotic analyses demonstrate that, regardless of the opponent, the tail of *Doedicurus* was capable of delivering functionally decisive strikes in close-range encounters. Glyptodonts flourished for at least 19 million years and weathered major ecological transitions, in part through the evolution of extensive armour and, in some lineages, extreme tail weaponisation, underscoring the adaptative value of this structure within the clade. Ultimately, however, even such formidable defences proved insufficient against a novel and unprecedented selective pressure: human predation (*52, 59, 60*).

Palaeorobotic physical modelling offers an experimental route to interrogate extinct performance landscapes when extant analogues and fossils alone are insufficient. When the fossil’s geometry and mass distribution are embodied at scale, the resulting strike produces a concrete force-time history that enables experimental validation of theoretical estimates. Robophysical models (*sensu* Lauder (*61*)) are therefore now required as a comparative method in palaeobiomechanics by holding background variables constant while selectively modifying specific traits and quantifying their performance consequences (*2*). Palaeorobotics is essential for testing functional hypotheses regarding the dynamics and control of motion systems by addressing them as measurable, falsifiable predictions (*62, 63*). Such anatomy-informed reconstruction of motion profiles, measurement of dynamics, and experimental validation with increasingly soft robots (*64*) facilitates moving beyond theoretical considerations toward quantitative mapping of how evolutionary and developmental constraints delimit motion and impact performance across clades, thus refining hypotheses and complementing analytical modelling. Extending bioinspiration into deep time palaeo-bioinspiration (*1*) thus expands the design space beyond extant biodiversity and encourages interdisciplinary translation from fossil systems to engineered technologies.

## Supporting information

Movie S1 (Supporting Material 2) High Speed Videography Footage of Collision Impact

## Acknowledgments

For initial investigations and the provision of photographs of Argentinian specimens, we thank Zoe M. Christen at the Natural History Museum of Basel. For conducting pilot studies on robophysical modelling of tail strike kinematics, we thank Olivier Lonneux, supervised by A.J., A.J.I., T.M.S. and K.L.V. For valuable insights on glyptodont tail autonomy, we thank Nicole Ramstein, supervised by K.L.V., T.M.S., and M.R.S-V. For access to complete glyptodont tails, we thank Agustín G. Martinelli at the Museo Argentino de Ciencias Naturales “Bernardino Rivadavia”, Analia M. Forasiepi at the National Scientific and Technical Research Council (CONICET, Mendoza) and Laureano R. González Ruiz (CONICET, Esquel). We also thank Gabriel Aguirre-Fernández and Jorge Domingo Carrillo-Briceño for their help in accessing specimens from the Department of Paleontology at the University of Zurich. For the acquisition of the 3D model of *Doedicurus clavicaudatus* PIMUZ A/V 459, we thank Dylan Bastiaans at the Natural History Museum Maastricht and Gizeh Rangel de Lazaro at the University of London. We would also like to thank Olivia Plateau at the University of Bern and the Natural History Museum of Bern for her contribution to discussions on defence index and ancestral state reconstruction. For the overall palaeorobotics test rig and device development, we thank Terence Fontana at Engineering Sciences Department of Empa. For force plate and hydraulic motor set up of life-sized glyptodont tail model, we thank Alexander Stutz from Engineering Sciences Department of Empa. For carpentry and CNC milling, we thank Phison Sisuwan from Empa. We thank Ziyou Wu and Leo Duggan of Engineering Sciences Department of Empa for insights.

## Author contributions

Conceptualization: AJ, KLV Methodology: AJ, FA, BWA Investigation: FA, BWA, BWE, AJI, AJ Formal analysis: KLV, AJ

Supervision: AJ, KLV, AJI, MRSV, TMS Writing – original draft: KLV, FA, AJ

Writing – review & editing: KLV, AJ, FA, BWA, BWE, AJI, MRSV, TMS

## Data, code, and materials availability

All data are available in the main text or the supplementary materials. The 3D models of the glyptodont tails used here for analyses, PIMUZ A/V 459, and for illustrative purposes, MACN-Pv 1780, MACN-Pv 9630, MACN-Pv 14998 and MACN-Pv 7126, are available from the open access online archive MorphoSource (Project ID: 000809881). No code is associated with the present study.

## Supplementary Materials

**Supplementary Material 1:** Materials and Methods

Figs. S1 to S4 Tables S1 to S2

**Supplementary Material 2:** Movie S1

Data S1 to S2

Supplementary Materials for

## Supplementary Material 1

Materials and Methods (N.B. all references included in main text file. References (*65–83*) only appear in the Materials and Methods section)

Figs. S1 to S4 Tables S1 to S2

## Supplementary Material 2

Movie S1 Data S1 to S2

## Materials and Methods

### Defence index generation and sample

Anatomy represents the empirical data on which our study is founded. To determine the type of defence for glyptodont tails, we extracted 25 tail morphological characters (71 to 95) from the 95 traditional characters of the most recent morphological phylogenetic matrix for glyptodonts (*25*), including 33 species that lived during the Neogene and Quaternary periods (fig. S2). Among them, 18 characters were selected to convert character states into ordered defence scores. Seven characters were deleted as they did not allow this conversion (characters 73, 74, 79, 80, 81, 90, and 91). One new defensive feature was defined to indicate tail composition, corresponding to the merge between characters 73 and 74 of the original matrix, resulting in 19 final defensive features. Due to missing data, uncertainty about tail composition, and the absence of a close sister species, the taxon *Parapropalaeohoplophorus septentrionalis* was removed from the analyses to minimize extrapolation. Conversion of phylogenetic character states into ordered defence scores follows three fundamental rationales described in the main text (see Data S1). Defence scores are thus ordered from the most protective to the most offensive for each trait. Some scores are redundant and correlated between defensive features. For example, the presence of a terminal tubercle necessarily forces the scoring of other defensive features. Here, the defensive features are not phylogenetic characters but inspired by a phylogenetic matrix. Consequently, these correlations serve as an optimization and do not represent a methodological obstacle to the generation of a defence index. To resolve the issues of gaps in the original phylogenetic matrix, the states for scoring have been reworked. For example, a caudal tube character in the original phylogeny has a score of 0 in the defensive feature, corresponding to the presence of a terminal tubercle, and therefore to the absence of a caudal tube. This approach, like the composite coding of Wilkinson (*65*), was used to eliminate all gaps. For missing data, we optimised the scoring according to the nearest sister taxon (*i.e.*, most parsimonious) and added to the matrix of Núñez-Blasco *et al*. (*25*) the specimens available at the Department of Paleontology at the University of Zurich, *i.e.*, *Glyptodon munizi* (PIMUZ A/V 463); *Glyptodon reticulatus* (PIMUZ A/V 4122), *Doedicurus clavicaudatus* (PIMUZ A/V 459 – the key specimen of this study), *Neosclerocalyptus* sp. (PIMUZ A/V 450 & 454), and *Panochthus tuberculatus* (PIMUZ A/V 434). The defence index for one species is computed as the sum of all scores divided by the number of defensive features. Defensive feature definitions and scores are available in data S1 and data S2, including defence index for each taxon and an indication of the scores optimized to overcome missing data. An anatomical plate is also presented in fig. S1. The central specimen of the study, *Doedicurus clavicaudatus* PIMUZ A/V 459, as well as four specimens from the palaeontological collections of the Museo Argentino de Ciencias Naturales ‘Bernardino Rivadavia’ (Buenos Aires, Argentina) for illustrative purposes (*Glyptodon reticulatus*, MACN-Pv 1780; *Neosclerocalyptus* sp., MACN-Pv 9630; *Panochthus* sp., MACN-Pv 14998; *Plohophorus* sp., MACN-Pv 7126), were scanned using structured light surface scanners, Artec EVA and Spider, in order to extract their morphology virtually.

### Physical reconstruction and test rig

The experimental series described in Figs. 2 and 3 were realised using an artificial glyptodont tail exhibiting similar weight and stiffness properties to the real glyptodont tail. The test setup to simulate a simplified glyptodont tail strike was built up on a multipurpose test bed (8 x 4 x 3.1 m) of the mechanical testing Laboratory of Empa, see Fig. 2 and movie S1. The impact tests of the tail strike were performed using a 6000 Nm/100° hydraulic rotary actuator of type Olear HYD-RO-AC HS010 2V OIL (Micromatic, USA, Log No. 117-28.570), so-called swivel motor, equipped with an angular position encoder of type RVDT R3D. The test controller Inova V. 1.13 (Inova, Czech Republic) was used to define the rotational motion of the tail. The artificial glyptodont tail consisted of a steel beam (total length: 2400 mm, inside length: 400 mm, cross-section: 30 x 60 mm) and a club tail made of plywood (length: 1000 mm, cross-section: 210 x 210 mm). These components were connected using two M10 screws, see Fig. 2. At the end of the club tail, a block of half circular section (radius: 105 mm) made of a bone substitute material (PU foam 62.5 PCF, Synbone, Switzerland), used as an impactor, was fitted with twin-sided adhesive tape. The total weight of the tail, combined with the extra weight, is about 50 kg. The hitting plate consists of a rectangular PU foam block (size: 180 x 130 mm, thickness: 3 mm PCF40 + 40 mm PCF20, Sawbones, USA), which was fixed with a twin-sided adhesive tape on a wood block (size: 250 x 250 mm, thickness: 160 mm). A polyester fibre mat was glued on the PU foam block. The whole construct was fixed with an elastic rope to the 350 kN load cell of type 737A (Erichsen, Germany, Log No. 125-12.068), which again was screwed with the vertical wall of the multipurpose test bed. Due to the strong expected strike, the sampling frequency of the recorded force signal was set to 10 kHz. The global view of the experiment was recorded by a Hero7 Black camera (GoPro, USA). Before the test, the tail was positioned at an initial angle of −45°±1°. The angle of the impactor when touching the PU foam block was defined as 20°. Different conditions/parameters were used, such as the hitting plate material, the impactor material, and the definition of the angular acceleration of the actuator. A total of five different test series were performed. In each test series, the targeted angular velocity was stepwise increased from 100°/s to 220°/s, thus reaching a velocity of the impactor just before touching the hitting plate of up to 4.3 m*s⁻¹ (measured with the high-speed camera of type MotionXtra HG-100K - Redlake, USA), the recorded rate was set to 1315 fps. The actuator is actuated backwards just before touching the hitting plate to avoid damaging the structure. Due to the relatively slow time response of the hydraulic actuator, the beam keeps rotating until it hits the hitting plate. Then, it bounces back, helping the actuator to accelerate in the opposite direction. The intent of this control strategy is to have an impact as short as possible, trying to reduce as much as possible the amount of force applied to the plate once after the impact. The impact velocity is evaluated offline by differentiating and filtering the measured angular velocity of the beam. The filter is realised in Matlab by means of a minimum order Infinite Impulse Response elliptic filter with a cutoff frequency of 10 Hz, allowing for zero-phase signal filtering (no delay is introduced in the velocity data). The controller actuator is not fast enough to track precisely the desired impact velocity; hence, a higher target velocity is fed to the controller. The desired impact velocity is obtained by trial, and error by increasing the target velocity experimentally. It has been observed experimentally that increasing the target speed above 270°/s would not cause a meaningful increment in the measured angular velocity of the beam. Consequently, this value was set as the upper limit of the impact velocity. Obtaining impact velocities with a precise and constant linear increment proved to be challenging; hence, the obtained increment is close to but not exactly 1 m·s⁻¹. In the reported experiments, the hydraulic actuator setup was configured as in table S1. To evaluate the mechanical performance of a biomimetic robot tail modelled after the tail of a glyptodont, we conducted a comprehensive finite element analysis (FEA) using explicit dynamics in ANSYS. The primary objective was to simulate the mechanical impact of both the robotic tail and the glyptodont fossil tail against a fixed force plate and compare their performance in terms of impact momentum and impulse generation. The geometric modelling involved creating detailed and accurate representations of both the bionic robot and glyptodont fossil tails. The dimensions and shapes were precisely matched to those of the glyptodont tail, ensuring that the models would provide a realistic basis for comparison. The geometric parameters are illustrated in fig. S4A. Assigning appropriate material properties to the components of both tails was crucial for achieving accurate simulation results. The properties of each material used in the analysis are listed in table S2. These properties were chosen based on their similarity to the natural and synthetic materials that would be used in the actual construction of the tails. The collision scenarios were set up to mimic real-world impact conditions. Both tails were simulated to collide with a fixed force plate at various velocities. fig. S4B shows the setup of these impact scenarios. The analysis included multiple impact velocities to capture a range of potential real-world conditions. During the simulations, the rotational impacts of the tails were analysed at different initial velocities (without an external force/torque). The explicit dynamics approach in ANSYS allowed for a detailed examination of the force-time relationship during each collision. The forces exerted by the tails on the force plate were recorded over time, and these data were used to calculate the impulse generated by each impact. The peak impact forces obtained in our experiment were then compared (Fig. 4) to those available for extant animals and human biomechanics (*4, 66–68*).

### Phylogeny and ancestral reconstruction

To address the evolution of the tail in glyptodonts, it is imperative to define a clear phylogenetic context. Because we removed a taxon from the matrix (see data S1 and sample), it was necessary to re-run the phylogenetic analysis of Núñez-Blasco *et al*. (*25*). Using the same matrix, apart from the exclusion of landmarks, we performed a maximum parsimony cladistic analysis using a heuristic search in PAUP v.4.0 (*69*). All the default parameters were respected by following the original analysis. We favoured convergences (DELTRAN) in the choice of our optimisation because they are assumed to be more probable (*70*). The strict consensus obtained differed slightly from the topology of Núñez-Blasco *et al*. (*25*) with the appearance of some polytomies. We therefore applied a backbone constraint to recover the topology of Núñez-Blasco *et al*. (*25*) by constraining the monophyly of the Santacruzian glyptodonts, the position of *Palaehoplophoroides* and *Palaehoplophorus*, the dichotomy within *Pseudoplophorus*, and the position of *Neosclerocalyptus pseudornatus* within the genus. The final topology obtained was then calibrated in time using the First Appearance Datum, favoured because of the interest in the appearance of the morphological traits of the tail. The time-calibrated tree is shown in fig. S2. This analysis is used here to update the topology of the phylogenetic tree for the examination we are carrying out in our study. The analysis is therefore not a reworking of the matrix of Núñez-Blasco *et al*. (*25*) to which we refer the reader for analyses focused on the phylogeny of glyptodonts. Using the time-calibrated tree and the defence index for each species, we performed the reconstruction of the ancestral defence indexes using maximum likelihood under Brownian motion with the *contMap* function of the *phytools* package (*71*) in R software (*72*).

### Biotic and abiotic data

To place the results obtained for the defence index and the case of *Doedicurus* in a broader evolutionary context, we integrated four selected biotic and abiotic data that may be associated with the evolution of glyptodonts in South America (*48*). For the temporal framework, we used the First Appearance Datum (FAD) and Last Appearance Datum (LAD) of each taxon (for a complete list of references, please refer to the study of Núñez-Blasco *et al*. (*25*). The stratigraphic extension was used for the phylogeny in fig. S2. However, we only used FADs for the calibration in the ancestral state reconstruction (Fig. 4) because we were interested in the appearance of defence acquisition rather than disappearance. For the variation in body mass, we used a compilation of different studies estimating body mass on the basis of the femur, skull or dental row (*46, 59, 73–83*). When no data was available for a given taxon, we used sister-group values if the proportions of comparable known material between the two taxa did not show aberrant size variations. Having estimates from different equations within the sampling is sub-optimal and constrained by the quality of the fossil record. Nevertheless, the ranges of variations are sufficiently informative to show the large variations in body mass within the clade. For the specific diversity of predators, we extracted the diversity of four major predator groups from Prevosti *et al*. (*20*): Carnivora (placental predators); Sparassodonta (marsupial predators); Phorusracidae (terror birds), and Sebecidae (terrestrial crocodilians). We intentionally ignored the Madtsoiidae (constrictor snakes) studied in Prevosti *et al*. (*20*) because they disappeared during the Early Eocene in South America. We preferred the study by Prevosti *et al*. (*20*) to more recent assessments to standardise the counting of the diversity of each predator group. For palaeoclimatic variation, we compared the curve based on benthic foraminifera δ^18^O values with the mean temperature difference relative to today based on Westerhold *et al*. (*37*). As several major environmental events are associated with paleoclimate variation, relating all the above variables to paleoclimate allows us to place the evolutionary history of glyptodonts within a broad environmental framework.

**Fig. S1.**
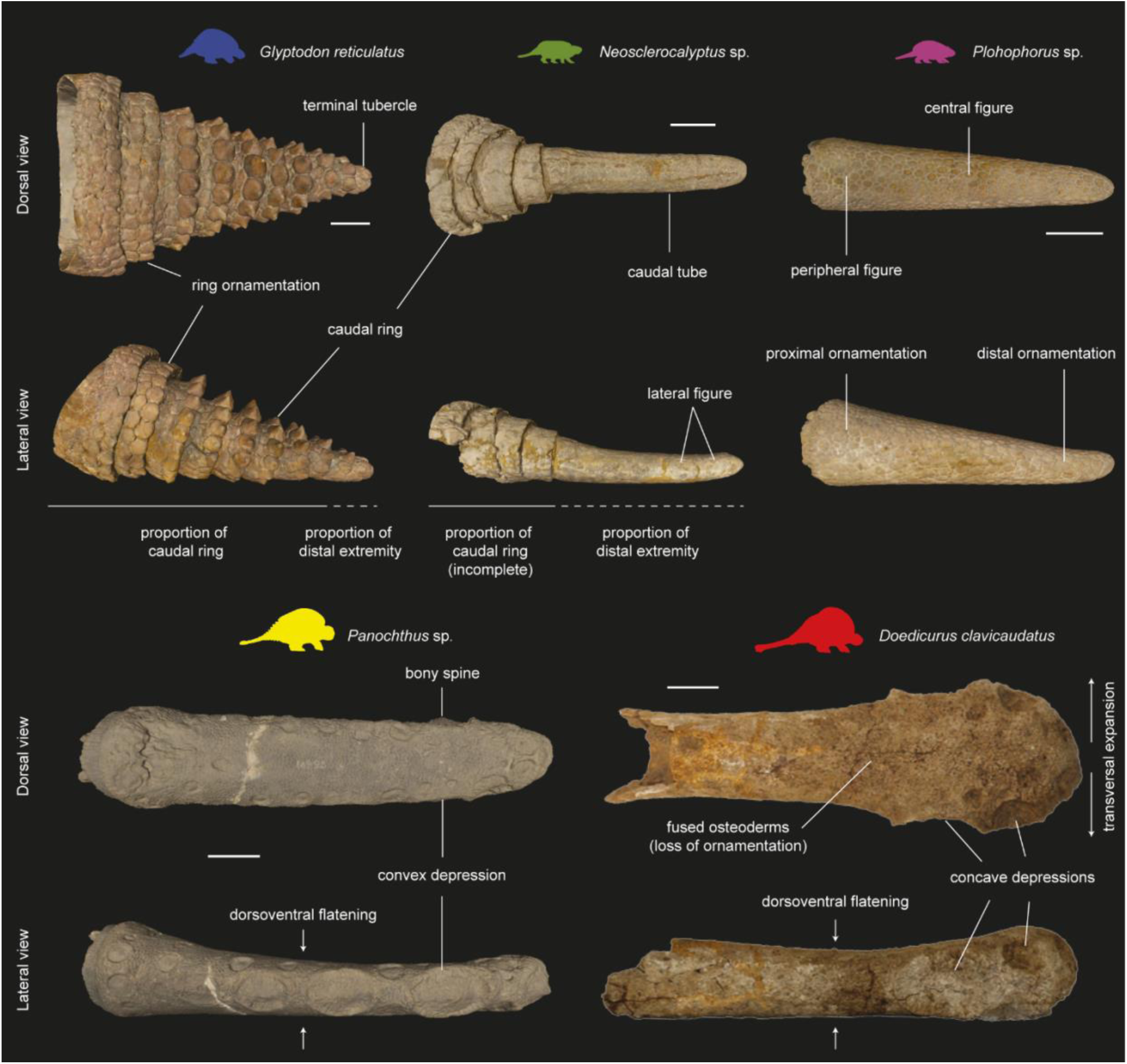
Tail anatomy and diversity. Illustration of the morphological characteristics used to convert phylogenetic characters into defence-related traits for the computation of the defence index. Specimens illustrated: *Doedicurus clavicaudatus*, PIMUZ A/V 459; *Glyptodon reticulatus*, MACN-Pv 1780; *Neosclerocalyptus* sp., MACN-Pv 9630; *Panochthus* sp., MACN-Pv 14998; *Plohophorus* sp., MACN-Pv 7126. The silhouettes are handmade. Scale bars, 10 cm.

**Fig. S2.**
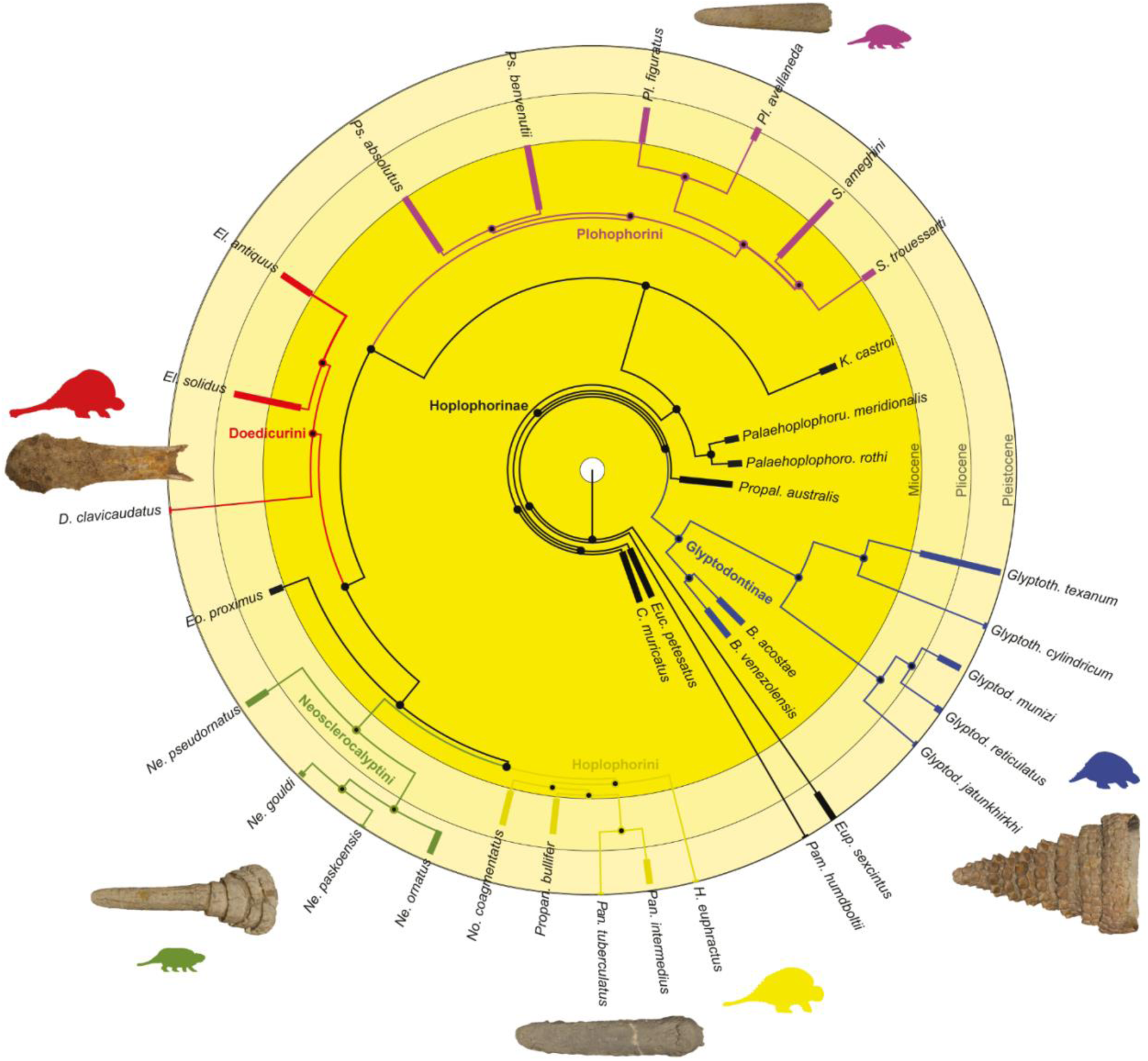
Glyptodont phylogeny. Time-calibrated phylogeny based on the character matrix of Núñez-Blasco *et al*. (*25*). Colour coding and stratigraphic ranges follow the original study. Illustrated glyptodont tails (not at scale) correspond to those shown in Fig. S1, presented here in dorsal view. The silhouettes are handmade.

**Fig. S3.**
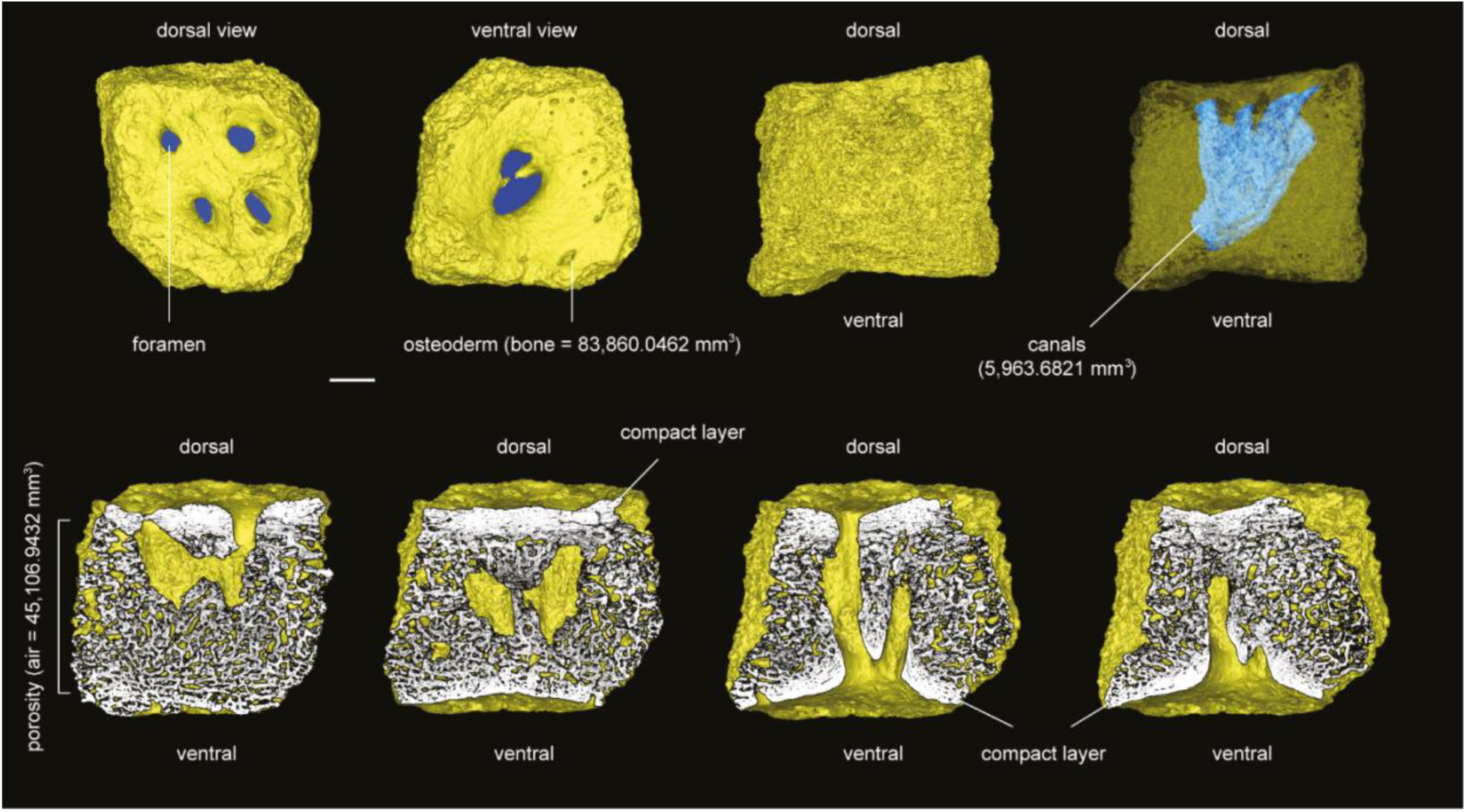
Osteoderm histology. Porosity (empty space) and vascularisation (blue canals) of an osteoderm (yellow 3D model) of the dorsal carapace of PIMUZ A/V 4106 *Doedicurus clavicaudatus*. Scale bar, 5 cm.

**Fig. S4.**
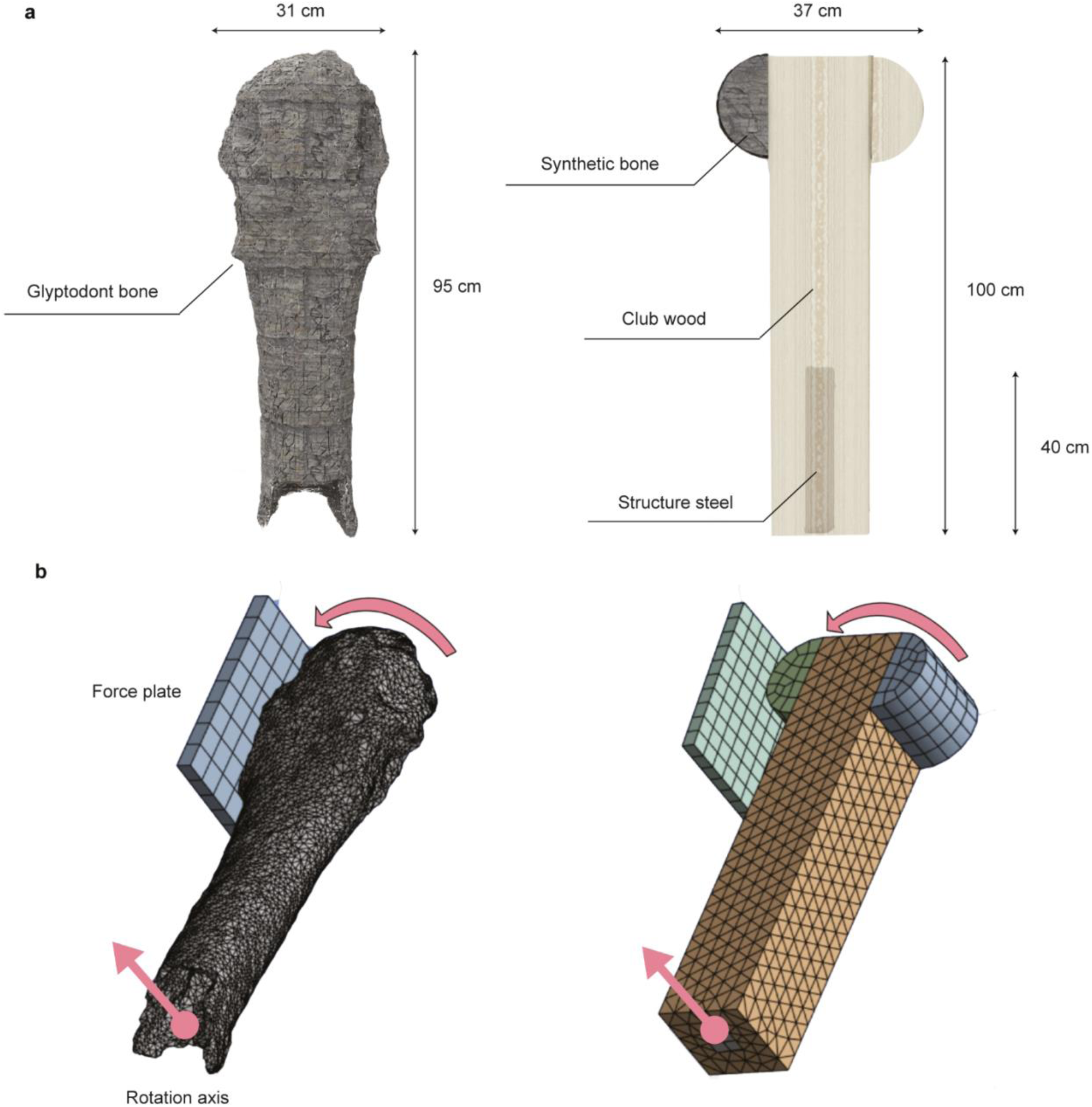
Palaeorobotic and glyptodont tails. Through FEA, we simulated the situation where the Palaeorobotic Tail and the Glyptodont Tail collide with the test force platform. (**A**) The size parameters of the Bionic Robot Tail and the Glyptodont Tail. (**B**) The collision scenarios of the two tails in finite element analysis.

**Table S1.** Details on the hydraulic actuator setup.

| Target controller angular velocity (deg/s) | SAA (deg)<sup>a</sup> | EAA (deg)<sup>b</sup> | Measured linear impact velocity (m/s) | Kinetic Energy (J)<sup>c</sup> |
| --- | --- | --- | --- | --- |
| 70 | -45 | 15 | 1.6 | 51.2 |
| 100 | -45 | 15 | 2.3 | 105.8 |
| 150 | -45 | 15 | 4.0 | 320.0 |
| 220 | -45 | 15 | 5.7 | 649.8 |
| 270 | -45 | 15 | 6.4 | 819.2 |

<sup>a</sup>SAA, start actuation angle; <sup>b</sup>EAA, end actuation angle; <sup>c</sup>kinetic energy considering a displacement of 1 cm.

**Table S2.** Material properties used in analyses.

| Name | Density (kg/m3) | Young's modulus (Pa) | Poisson's ratio |
| --- | --- | --- | --- |
| Glyptodont bone | 1.40E+03 | 2.00E+10 | 0.40 |
| Synthetic bone | 1.00E+03 | 2.00E+10 | 0.40 |
| Club wood | 6.00E+2 | 1.00E+09 | 0.30 |
| Structure steel | 7.65E+03 | 2.00E+11 | 0.30 |

### Movie S1. (separate file)

High-speed video footage of the life-sized palaeo-inspired robophysical model of *Doedicurus* tail strike impacts. Zoomed in view of contact zone on impact of high-density distal tail club (Synbone) striking a target force plate with porous osteoderm inspired element; shown 50 times slowed (recorded at 1315 fps from top view). Bird eye view of life-sized robophysical model of hydraulic motor moving tail club towards force plate across tested velocities of 1 to 5 m·s⁻¹, playing in real time, followed by zoomed in view of tail club striking a force plate, respectively.

### Data S1. (separate file)

Coding strategy for morphological characters used in the computation of the defence index.

### Data S2. (separate file)

Scoring from phylogenetic characters (K) from (*25*), indexes, First Appearance Datum (FAD), Last Appearance Datum (LAD) and Body Mass (BM) for each species considered in the present study. The shaded numbers correspond to scores estimated based on sister groups.

