## Supplementary File S1 for "PALAEOROBOTICS UNCOVERS DEVASTATING MAMMALIAN TAIL STRIKE DYNAMICS"

**Data S1: Defence index generation**

The defence index is based on the conversion of the morphological characters (= K) of the tail from one of the most recent phylogenies of glyptodonts^25^ into defensive coding. The phylogenetic matrix contains 25 Ks focused on the tail. After an initial evaluation, 18 Ks were converted, two were merged to create a new K with defensive coding, and the last 5 were deleted as not suitable for conversion (*i.e.*, not informative or following any rational for conversion). Considering the tail as a means of engaging in a strike, the conversion of Ks is based on simple rationales:

1) a long caudal tube is considered more optimal for striking than a short caudal tube or terminal tubercle.

2) a tail better adapted to striking is supposed to show less ornamentation for improved robustness but high vascularity for bone remodelling and other tissues.

3) any structure that increases the complexity of the tail shape and surface (excluding ornamentation) of the distal end is considered as weaponisation.

Following these rationales, the states of each K are transformed into scores and ordered in a manner such that the score considered to be the most protective corresponds to the lowest value (*i.e.*, 1) and the score considered to be the most offensive corresponds to the highest value (*i.e.*, up to 7). This coding enables us to define minimum and maximum defence values ranging from the most protective to the most offensive within each K. For a given species, the sum of the scores divided by the number of Ks (n=19) enables us to define a comparable defence index between species. This approach allows us to incorporate as many species as possible despite very different morphologies and the missing data inherent in fossil records, here optimised according to the sister group in each case. Unlike phylogenetic studies, it should be noted that the redundancy of a morphological trait in the matrix (*e.g.*, terminal tubercle vs. caudal tube) was preserved to favour the polarisation of defence scores. The list of all the Ks is given below with the associated references. Changes from the initial K of Núñez-Blasco *et al*.^25^ are also indicated. The coding and calculation of the defence index for each species is available in Table S1. For examples of anatomical variation, please refer to Extended Data Fig. 1.

**K1 – number 71 in Núñez-Blasco *et al*. (*25*)** **– Morphology of the exposed surface of the osteoderms of the caudal rings:**

**Score 1:** distal series of osteoderms with tuberculate surface (state 2 in Núñez-Blasco *et al*. (*25*));

**Score 2:** flat and without ornamentation or with peripheral figures (states 0 and 1 in Núñez-Blasco *et al*. (*25*));

**Score 3:** smooth with foramina (state 3 in Núñez-Blasco *et al*. (*25*)).

**K2 – number 72 in Núñez-Blasco *et al*. (*25*)** **– Peripheral figures on the exposed surface of the distal series of the caudal ring osteoderms:**

**Score 1:** absent (state 0 in Núñez-Blasco *et al*. (*25*));

**Score 2:** present (state 1 in Núñez-Blasco *et al*. (*25*)).

**K3 – fusion between number 73 and 74 in Núñez-Blasco *et al*. (*25*)** **– Caudal armour composed of:**

**Score 1:** caudal rings and a terminal tubercle;

**Score 2:** only caudal rings without specific distal structure;

**Score 3:** caudal rings and a caudal tube.

**K4 – number 75 in Núñez-Blasco *et al*. (*25*)** **– Percentage occupied by caudal rings in the caudal armour:**

**Score 1:** rings representing more than 90% of the total length (state 0 in Núñez-Blasco *et al*. (*25*));

**Score 2:** rings representing 80% of total length (state 1 in Núñez-Blasco *et al*. (*25*));

**Score 3:** rings representing up to 60% or less of total length (state 2 in Núñez-Blasco *et al*. (*25*)).

**K5 – number 76 in Núñez-Blasco *et al*. (*25*)** **– Fusion degree of the caudal rings composing the caudal tube:**

**Score 1:** no caudal tube;

**Score 2:** without developing a complete tube (state 0 in Núñez-Blasco *et al*. (*25*));

**Score 3:** developing a complete short tube with very visible rings (state 1 in Núñez-Blasco *et al*. (*25*));

**Score 4:** developing a complete long tube with very visible rings (state 2 in Núñez-Blasco *et al*. (*25*));

**Score 5**: developing a complete long tube and rings still visible (state 3 in Núñez-Blasco *et al*. (*25*));

**Score 6**: developing a complete long tube without visible rings (state 4 in Núñez-Blasco *et al*. (*25*)).

**K6 – number 77 in Núñez-Blasco *et al*. (*25*)** **– Transverse section of distal third of caudal tube:**

**Score 1**: no caudal tube or complete tube;

**Score 2**: nearly circular (state 0 in Núñez-Blasco *et al*. (*25*));

**Score 3**: dorsoventrally flattened (state 1 in Núñez-Blasco *et al*. (*25*)).

**K7 – number 78 in Núñez-Blasco *et al*. (*25*)** **– Sutures between the fused rings of the caudal tube:**

**Score 1:** no caudal tube or complete tube;

**Score 2:** visible (state 0 in Núñez-Blasco *et al*. (*25*));

**Score 3:** not visible (state 1 in Núñez-Blasco *et al*. (*25*)).

**K8 – number 82 in Núñez-Blasco *et al*. (*25*)** **– Osteoderms of proximal portion of caudal tube (dorsal view):**

**Score 1:** no caudal tube or peripheral figures absent (state 0 in Núñez-Blasco *et al*. (*25*));

**Score 2:** rosette pattern with one row of peripheral figures around central one poorly developed, not completely surrounding central figures (states 1-3 in Núñez-Blasco *et al*. (*25*));

**Score 3:** presence of more than one row surrounding central figure (state 4 in Núñez-Blasco *et al*. (*25*));

**Score 4:** no discernible ornamentation due to osteoderm fusion.

**K9 – number 83 in Núñez-Blasco *et al*. (*25*)** **– Osteoderms of distal portion of caudal tube (dorsal view):**

**Score 1:** no caudal tube or peripheral figures absent (state 0 in Núñez-Blasco *et al*. (*25*));

**Score 2:** rosette pattern with one row of peripheral figures around central one poorly developed, not completely surrounding central figures (states 1-3 in Núñez-Blasco *et al*. (*25*));

**Score 3:** presence of more than one row surrounding central figure (state 4 in Núñez-Blasco *et al*. (*25*));

**Score 4:** no discernible ornamentation due to osteoderm fusion.

**K10 – number 84 in Núñez-Blasco *et al*. (*25*)** **– Ornamentation pattern of the dorsal osteoderms of the caudal tube:**

**Score 1:** without ornamentation or no complete caudal tube (state 0 in Núñez-Blasco *et al*. (*25*));

**Score 2:** rosette pattern (state 1 in Núñez-Blasco *et al*. (*25*));

**Score 3:** reticular pattern (state 2 in Núñez-Blasco *et al*. (*25*));

**Score 4:** smooth surface with numerous foramina (state 3 in Núñez-Blasco *et al*. (*25*)).

**K11 – number 85 in Núñez-Blasco *et al*. (*25*)** **– Peripheral figures of the caudal tube with polygonal morphology, with straight edges:**

**Score 1:** no caudal tube or complete tube;

**Score 2:** absent (state 0 in Núñez-Blasco *et al*. (*25*));

**Score 3:** present (state 1 in Núñez-Blasco *et al*. (*25*)).

**K12 – number 86 in Núñez-Blasco *et al*. (*25*)** **– Number of lateral figures of the caudal tube:**

**Score 1:** no caudal tube or complete tube.

**Score 2:** no differentiated lateral figures (state 4 in Núñez-Blasco *et al*. (*25*));

**Score 3:** four or fewer lateral smooth figures (state 1 in Núñez-Blasco *et al*. (*25*));

**Score 4:** five or more lateral smooth figures (state 0 in Núñez-Blasco *et al*. (*25*));

**Score 5:** convex insertion structures (state 2 in Núñez-Blasco *et al*. (*25*));

**Score 6:** concave insertion structures (state 3 in Núñez-Blasco *et al*. (*25*)).

**K13 – number 87 in Núñez-Blasco *et al*. (*25*)** **– Lateral figures of the caudal tube:**

**Score 1:** no caudal tube or complete tube.

**Score 2:** without peripheral figures (state 0 in Núñez-Blasco *et al*. (*25*));

**Score 3:** with few peripheral figures, they do not surround the entire figure (state 1 in Núñez-Blasco *et al*. (*25*));

**Score 4:** the peripheral figures surround the dorsal and ventral margins (state 2 in Núñez-Blasco *et al*. (*25*));

**Score 5:** the peripheral figures surround the entire central figure (state 3 in Núñez-Blasco *et al*. (*25*));

**Score 6:** the peripheral figures surround the entire central figure and there are also several rows around the central figure (state 4 in Núñez-Blasco *et al*. (*25*));

**Score 7:** very large lateral figures without peripheral figures discernible due to osteoderm fusion.

**K14 – number 88 in Núñez-Blasco *et al*. (*25*)** **– General characteristics of the distal portion of caudal armour:**

**Score 1:** no caudal tube or terminal tubercle;

**Score 2:** Terminal tubercle composed of two or three transverse rows of ankylosed osteoderms (state 3 in Núñez-Blasco *et al*. (*25*));

**Score 3:** Caudal tube composed of four or five transverse rows of osteoderms with observable sutures and lacking peripheral figures (state 0 in Núñez-Blasco *et al*. (*25*));

**Score 4:** Cylindrical-conical caudal tube composed of ankylosed osteoderms, with five to seven large lateral figures (state 1 in Núñez-Blasco *et al*. (*25*));

**Score 5:** Cylindrical-conical caudal tube composed of ankylosed osteoderms with large lateral depressions; (state 2 in Núñez-Blasco *et al*. (*25*)).

**K15 – number 89 in Núñez-Blasco *et al*. (*25*)** **– Characteristics of the exposed surface of the lateral figures of the caudal tube:**

**Score 1:** no caudal tube or complete tube;

**Score 2:** smooth surface (state 0 in Núñez-Blasco *et al*. (*25*));

**Score 3:** rough surface with a convex centre (state 2 in Núñez-Blasco *et al*. (*25*));

**Score 4:** rough surface with a concave centre (state 1 in Núñez-Blasco *et al*. (*25*)).

**K16 – number 92 in Núñez-Blasco *et al*. (*25*)** **– Terminal tubercle of caudal armour:**

**Score 1:** no caudal tube or terminal tubercle;

**Score 2:** formed by three fused rings (state 1 in Núñez-Blasco *et al*. (*25*));

**Score 3:** formed by two fused rings (state 0 in Núñez-Blasco *et al*. (*25*));

**Score 4:** No terminal tubercle but a caudal tube.

**K17 – number 93 in Núñez-Blasco *et al*. (*25*)** **– Large lateral figures at the apex of the caudal tube:**

**Score 1:** no caudal tube;

**Score 2:** present (state 0 in Núñez-Blasco *et al*. (*25*));

**Score 3:** absent (state 1 in Núñez-Blasco *et al*. (*25*));

**Score 4:** lateral figures replaced by convex or concave depressions.

**K18 – number 94 in Núñez-Blasco *et al*. (*25*)** **– Morphology of the terminal part of the caudal armour:**

**Score 1:** no caudal tube or complete tube.

**Score 2:** not transversely expanded (state 0 in Núñez-Blasco *et al*. (*25*));

**Score 3:** slightly transversely expanded (state 1 in Núñez-Blasco *et al*. (*25*));

**Score 4:** very transversely expanded (state 2 in Núñez-Blasco *et al*. (*25*)).

**K19 – number 95 in Núñez-Blasco *et al*. (*25*)** **– Ornamentation of the proximal-ventral region of the caudal tube, with accessory figures between rows two and three of osteoderms:**

**Score 1:** no caudal tube or complete tube;

**Score 2:** numerous and contiguous (state 0 in Núñez-Blasco *et al*. (*25*));

**Score 3:** isolated (state 1 in Núñez-Blasco *et al*. (*25*));

**Score 4:** absent (state 2 in Núñez-Blasco *et al*. (*25*));

**Score 5:** no discernible ornamentation due to osteoderm fusion.
